# Tracing the influence of land-use change on water quality and coral reefs using a Bayesian model

**DOI:** 10.1101/112250

**Authors:** Christopher J. Brown, Stacy D. Jupiter, Simon Albert, Carissa J. Klein, Sangeeta Mangubhai, Maina Mbui, Peter Mumby, Jon Olley, Ben Stewart-Koster, Vivitskaia Tulloch, Amelia Wenger

## Abstract

Coastal ecosystems can be degraded by poor water quality. Tracing the causes of poor water quality back to land-use change is necessary to target catchment management for coastal zone management. However, existing models for tracing the sources of pollution require extensive data-sets which are not available for many of the world’s coral reef regions that may have severe water quality issues. Here we develop a hierarchical Bayesian model that uses freely available satellite data to infer the connection between land-uses in catchments and water clarity in coastal oceans. We apply the model to estimate the influence of land-use change on water clarity in Fiji. We tested the model’s predictions against underwater surveys, finding that predictions of poor water quality are consistent with observations of high siltation and low coverage of sediment-sensitive coral genera. The model thus provides a means to link land-use change to declines in coastal water quality.

## Introduction

The management of activities on land to avoid pollution run-off to the ocean is important for the conservation of many coastal marine ecosystems ^1–3^. Deforestation and farming increase run-off of nutrients and sediment that can flow out to the ocean ^4^. In the ocean, sediment and nutrient pollutants can decrease water clarity and shade or smother habitats, reducing diversity of benthic organisms ^5^, habitat complexity and fish diversity ^6,7^. Thus, in places where coastal waters are strongly influenced by freshwater run-off the management of marine ecosystems requires actions in connected terrestrial and freshwater habitats.

Management of run-off to coastal marine ecosystems requires identifying the source of impacts to ecosystems, so that appropriate actions can be taken to reduce threats. Between the land-use change and changes in marine ecosystems, multiple processes are operating across space and time that affect marine sediment concentrations: for instance, deforestation causes increased sedimentation in rivers and floodplains e.g. ^4,8,9^, rivers transport sediments to the ocean e.g. ^10^ and in the ocean sediments are dispersed to reefs e.g. ^11^. Given that the ocean can disperse sediments widely, water quality at a single location in the marine environment may be influenced by rivers that drain multiple catchments. Thus, attributing declines in coastal water quality to its cause on land is difficult, which hinders identifying priorities for management actions on land, like the best locations for re-vegetation ^4^.

The methods used to trace the source of water quality issues to their causes on land are generally data-intensive. For instance, on the Great Barrier reef, millions of dollars and years of research have been in invested to trace the source of poor water quality ^12^. Satellite remote sensing products, *in situ* water quality measurements and river discharge measurements have been used to trace the extent of influence of river flood plumes ^13^ and estimate potential improvements in water clarity from acting to reduce river sediment loads ^14^. However, the investment of time and resources required to implement these approaches is not feasible in many developing countries, where run-off can have severe negative impacts on the livelihoods of people that rely on coastal ecosystems ^15^. For instance, people in Fiji are reliant upon coral reefs for fisheries and tourism, an ecosystem that is threatened by land-use change ^16^. There is limited historical data, funding and capacity to undertake additional science to support a new government initiative for integrated coastal zone management. Capitalizing on political opportunities for coastal planning requires methods that can be implemented with existing and freely available data-sets ^17^.

Three approaches offer hope for data limited regions. The first uses soil erosion equations, such as in those in the INVEST toolbox ^18^, to inform land areas contributing the greatest sediment and nutrient loads to river mouths. The second approach relies on simple models of reef exposure to river flood plumes based on Geographic Information Systems analysis (GIS) e.g. ^11,19,20^. A weakness of these modeling approaches is that they are not quantitatively validated against local datasets, and parameters are estimated using expert opinion or extrapolated from other study areas, often with very different climates and soil conditions ^8^. A third approach, the analysis of satellite data for indices of water quality offers a way to obtain quantitative measurements even in data-limited regions e.g. ^13^. However, satellite measurements should be corrected locally for biases, for instance from benthic reflectance ^21^. Further, satellite measurements cannot be used on their own to trace the source of poor water quality back to land. An appropriate statistical modeling framework could integrate these three approaches, drawing strengths from each, is required.

Here we combined satellite measurements with GIS models and catchment models to resolve the contributions of different catchments to water quality, specifically turbidity, at coral reefs using a hierarchical Bayesian model. We applied our model to estimate catchment contributions to turbidity around Vanua Levu, Fiji, where information on sediment run-off is needed to inform coastal planning ^22^. Because field data are often poorly controlled and there are inherent errors in satellite measurements of water quality ^13,14^, we further tested our model under idealistic conditions to suggest further scope for improvement in the model and priorities for collecting empirical data. Our approach required freely available data on water quality and land-use (from satellite imagery) and rainfall, making it broadly applicable for linking catchments to water quality, even in data-limited regions, for use in integrated land-sea management plans.

## Methods

### Overview

We approach the problem of determining the contribution of different catchments to satellite measurements of turbidity in the ocean by developing a Bayesian hierarchical model. The model simultaneously estimates the dispersion of sediments from sources (e.g. river mouths) and the relative influence of different sources on ocean turbidity. The model was hierarchical because the influence parameters were scaled by independently derived estimates of sediment loadings. Sediment loadings themselves were calculated from catchment land-use and rainfall data using a simple model of sediment run-off. To test Bayesian model’s predictions of turbidity, we related model estimated turbidity values to observed benthic habitat data. Finally, we perform a power analysis where we test the Bayesian model’s ability to recover known parameter values for a simulated coastline.

### Bayesian model

The core of our model was a function that described the influence of different sediment sources on ocean turbidity at different distances from each source. In all equations below we use Greek letters to represent parameters that were estimated. The likelihood of the satellite observing turbidity value *y_i_* at location *i* was specified:

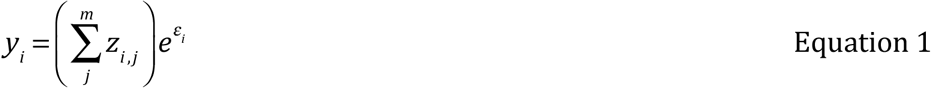

where the summation is over *m* sediment sources, *j*, εi are normally distributed error terms with precision τ_y_ and *Z_i,j_* was a latent variable representing the influence of source *j* (e.g. a river mouth) on ocean location *i*. We rescaled the turbidity measurements by subtracting the minimum value then adding a small number so that estimation of an intercept parameter was unnecessary ^23^.

The model described the declining influence of a source *j* on turbidity at an ocean site (*Z_i,j_*) using a power function:

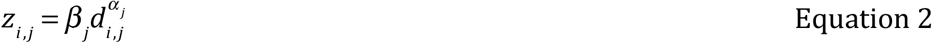

where β_*j*_ was the influence of source *j* on turbidity at a distance of zero (e.g. at a river mouth), α_*j*_ was a scaling parameter that controls the dispersion of sediment, and *d_i,j_* was a matrix of distances (in kilometres) from ocean sites to sources. The parameters α_*j*_ were expected to be negative if turbidity declines at greater distance from sources. We let the dispersion parameter vary by sources, however, in practice allowing each source to have a unique α would result in issues with parameter identification. Thus we suggest that α values are restricted to one or just a few values. We also tested the model by replacing equation 2 with an exponential function, which assumes diffusion of sediments across space. However, model fits from the exponential function were poor, so we proceeded with the power function.

We included a second hierarchical level in the model to allow for source influences (*β_j_*)to scale with estimates of sediment yield from catchments:

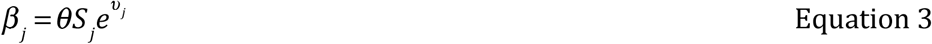

Where the parameter θ can take values >0 and rescaled estimates of sediment loadings coming from rivers, *S_j_,* into the units of the satellite measurement and υ_j_ were normally distributed errors with precision τ_υ_. Sediment loadings themselves are estimated according to land use and rainfall patterns within each catchment, outlined below.

The hierarchical level was an important strength of this modeling approach. The problem of attributing the contribution of multiple catchments to ocean turbidity is underdetermined, because there is no one unique solution. Constraining catchment influences to scale with their sediment loadings effectively requires the relative order of catchment influences to remain similar to the order of their sediment loadings. This constraint helps constrain the plausible parameter space.

We used vague priors for all parameters and priors were specified as follows: τ_y_, τ_υ_ and α_*j*_ had uninformative gamma priors, and θ had an uninformative log-normal prior (See appendix B for further details).

We used Markov-Chain-Monte-Carlo (MCMC) sampling to numerically simulate posterior distributions of the modeled parameters ^23^. To implement numerical simulations, we used JAGS version 3.3.0 ^24^, controlled from the R programming language ^25^, using the Coda package ^26^ to evaluate model fits (Supplementary Material Appendix A). For all model runs, we used the Gelman-Rubin statistic ^27^ to evaluate convergence and only accepted models where this statistic was <1.05 for all variables.

The model could be applied to different water quality variables (e.g. turbidity, salinity) over different time-scales, provided the estimates of catchment size and the water quality variables are measured at consistent time-scales. For instance, the model could be applied to a pulse event, such as a tropical storm, to estimate the contribution of different rivers to sediment pollution in the ocean. The model could also be applied to time-integrated measures of pollutant exposure e.g. ^19^, which is the approach we take here.

### Case-study

We estimated catchment influences on ocean turbidity in the waters around Vanua Levu, the second largest island of Fiji. Integrated Coastal Management (ICM) plans are currently being developed at the provincial level for Vanua Levu. In Vanua Levu, there is concern from government and communities that accelerating economic development on land, including mining agriculture, road building and forestry will impact fisheries, tourism and the ecological integrity of marine protected areas ^28^. To address these concerns, coastal communities in Vanua Levu’s Bua province are currently working with government and a non-governmental organization (The Wildlife Conservation Society, WCS) to design and implement an ICM plan based on national ICM frameworks for Fiji ^22^. The ICM plan will aim to balance terrestrial economies, marine economies and ecological health. Therefore, the ICM plan requires information on where development on land can have minimal impacts to marine ecosystems and economies. However, there are limited data on the influence of land-use change on coastal water quality to support this decision process and limited funding and time to support further data collection. Thus, the ICM process would benefit from rapid advice on where development may have the greatest impact on water quality.

### Data for Vanua Levu

We fitted the Bayesian model to remotely sensed turbidity measurements from the coastal waters around Vanua Levu. We used satellite data from the Medium Resolution Imaging Spectrometer (MERIS), part of the European Space Agency’s Envisat platform ^29^. We downloaded images for the Level 2 products for turbidity for the years 2003-2011 (data provided in Formazin Turbidity Units). The turbidity product has been empirically validated for other regions with similar water types ^13,30^. The satellite pixels were summarized onto a standardized grid of 402 by 402 metres, removing any pixels with a low quality reading (quality flag <0.01). We also masked pixels on reefs, in shallow water and all pixels next to reefs or shallow water to minimize the confounding influence of benthic reflectance ^21^. Reefs were identified using a global reef database ^31^.

Turbidity measurements were summarized as the geometric mean of values across all years in the wet season (119 images, November to April) and standardized by the mean value. Standardization was performed because turbidity has not been validated against local in situ data, and we were interested in the spatial patterns, not the absolute values. We also created summaries using the maximum, minimum and frequency of high (>2SD) turbidity events, however these all had similar spatial patterns, so we focus our analyses on the means summary. The summary was resampled to a resolution of 3.12km by 3.14 km, resulting in 1250 pixels with turbidity measurements in the study region. Resampling was performed because convergence of the MCMC algorithm was slow using the full resolution data. Resampling to a lower resolution preserved spatial patterns in turbidity and exploratory analysis indicated there was little bias in parameter estimates when resolution was reduced. The pixels that were not used in model fitting were retained for evaluating model fit. Model fit was evaluated using the residual mean square error. Parameters, priors and model code are provided in Supplementary Material Appendix B.

Sediment yield from each river mouth was estimated for each catchment as:

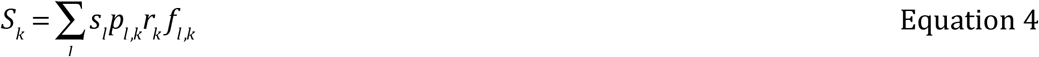

Where *S_k_* in mg is the summed product over the landuses *l*, *S_l_* is the sediment yield of a land-use (per mg/L of rainfall), *p_l,k_* is the proportion of rainfall that runs off a landuse (L^−1^), *r_k_* is the total rainfall (L) in a catchment in the wet-season and *f_l,k_* is the proportion of area under a landuse in a catchment (Σ *f _l,k_* = 1). The proportion of rainfall that ran off a given catchment increased with increases in that catchments spatially averaged wet season rainfall ^32^.

We consider two land-uses: forested versus deforested (including farmland, bare soil and settlements). There were 74 catchments in the study region and for model fitting we scale the *S_k_* relative to the largest catchment. Delineation of catchment boundaries and data to calculate source sediment contributions (Equation 4) were derived using freely available GIS products and programming routines (Supplementary Material Appendix C).

Catchments with river mouths within 5km of each other were aggregated together before fitting the Bayesian model. Aggregation was necessary because the influence of nearby river mouths on turbidity was not identifiable by the model. For aggregated river mouths distance from the rivers to each ocean pixel was taken as the mean distance from the river mouths. River sediment loadings, *S_k_* were summed over the groups to obtain 28 grouped river mouths, *S_j_*. Aggregation of river mouths meant we could only discriminate sediment contributions from groups of neighboring catchments, however simulating testing indicated that the aggregation procedure significantly reduced bias in parameter estimates (see below).

Initially we fit the model with a single dispersion parameter (α) for all sources. However, examination of model fits indicated that this model tended to under-predict turbidity on the north-west facing coast of Vanua Levu and over-predict turbidity on the south-east coast.

Winds in Fiji are predominantly easterlies ^33^, suggesting that sediment dispersion might vary on the two coasts. Therefore we re-fitted the model allowing unique dispersion parameters for southern and northern coasts. We report results from both models, including statistics for model selection: the predictive loss and the deviance information criteria ^23,24^. Lower values of predictive loss and deviance information criteria indicate a more parsimonious model.

### Impact of poor water quality on benthic habitats

We conducted an independent verification of water quality predictions by assessing whether the composition of benthic habitats was consistent with the Bayesian model’s predicted gradient in turbidity. We used mean turbidity as predicted by the model with two dispersion parameters. Surveys of coral reef benthic habitats were conducted by WCS at 168 sites around Vanua Levu. Point intercept surveys were performed using 2-6 replicates of 50 metre long transects at each site, recording benthic habitat categories at 0.5 metre intervals according to a standard classification adapted from Hill and Wilkinson ^34^.

For each site we calculated change in percent cover of three benthic habitat types that are most likely to respond to high turbidity: silt, sediment sensitive scleractinian coral genera (genera and justification are in Supplementary Material Appendix D) and algae. Silt cover and percent cover of algae (which included macro algae and cyanobacteria) were expected to increase with increased turbidity, cover of sediment sensitive coral genera was expected to decline with increased turbidity. For the three habitat types (silt, algae and coral), we fitted linear models to test for a relationship between predicted turbidity and the cover of each habitat. For each habitat type, we used linear models and transformations appropriate to the distribution of residuals (Supplementary Material Appendix D).

### Simulation testing

We used a simulation study to test the model’s ability to estimate source contributions to pollution accurately and precisely for different geographies and data types. We simulated a 100 km long linear coastline with three river mouths. The ocean environment extended out to 100 km from shore and contained 300 (30 by 10) ocean pixels. For each simulation test we used the power functional form (equation 2) to simulate 25 water quality data-sets with random measurement errors. We then fit equations 2–3 to each simulated data-set to see if it could recover the original parameters (see Supplementary Material Appendix E for details of the MCMC algorithm).

We were particularly interested in the model’s ability to predict turbidity at ocean pixels and estimate the contribution of each source to water quality. We used two statistics to evaluate the model’s performance to estimate the source contributions ^35^. The first was the mean relative error (MRE), which is a measure of bias in parameter estimation. The second was the mean coefficient of variation, which quantifies precision. We also evaluated bias in estimation of ocean turbidity using the residual mean square error comparing model predicted turbidity to simulation turbidity without observation error (Supplementary Material Appendix E).

Initially, we assumed the source contributions to turbidity were equal and they were position at 100km, 200km and 300km along the coastline. In the first simulation test, we varied the standard deviation of the observation error and ocean dispersal *a* (Supplementary Material Appendix E). Based on these initial tests, we fixed the error and *a* at 0.5 and 1.25 for further testing.

The second simulation test was to explore the model’s ability to partition the contribution of the sources to water quality. We then ran crossed trials where the southern-most source was iteratively moved towards the center of the coast. We also iteratively increased the magnitude of one source’s contribution to water quality.

## Results

### Case-study for Fiji

Satellite measurements of turbidity indicated a gradient in the geometric mean of wet-season turbidity with high values near to river mouths (Fig. 1). Values tended to be higher on the north-coast of Vanua Levu, where there has been extensive land-clearing.

**Fig 1.**
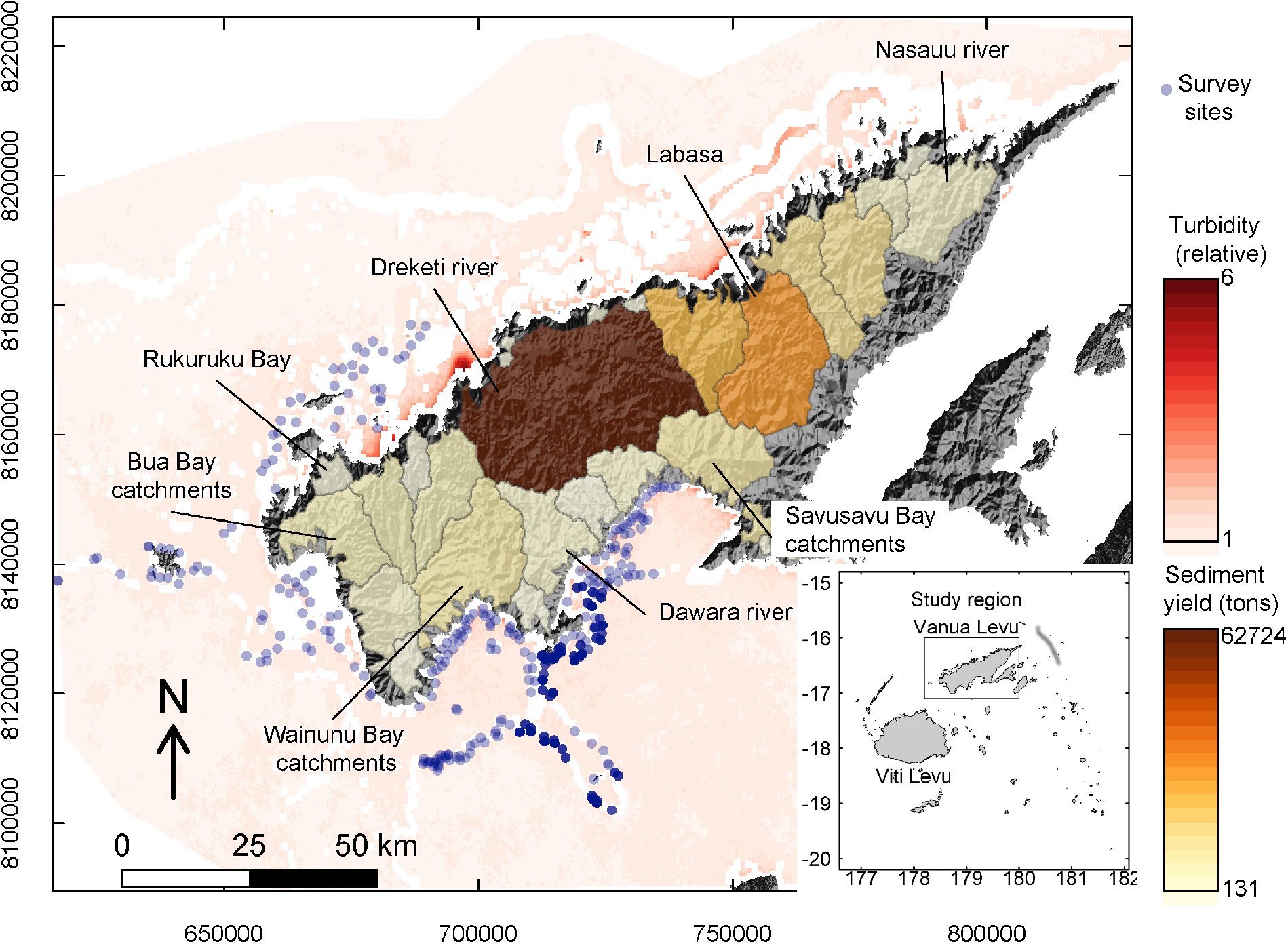
Map of the study region, showing grouped catchments (coloured polygons, overlaid on a hillshade map) with their estimated sediment yield (tons per wet season) and reef survey sites. White areas were excluded from analysis due to shallow water or being too far from the catchments. Ocean colour scale shows mean turbidity estimated from satellites. Inset shows the study region's location in relation to Fiji's largest islands.

Comparing the two model fits, the model with unique dispersion parameters for each coast provided a more accurate fit (lower root-mean square error) and was more parsimonious despite the extra parameter (lower DIC, Table 1). The estimates of the dispersion parameters also suggested that the dispersion parameter was significantly different on north and south coasts (compare overlap of 95% credibility intervals in Table 1). For further analysis we proceed with the two dispersion parameter model.

**Table 1.**
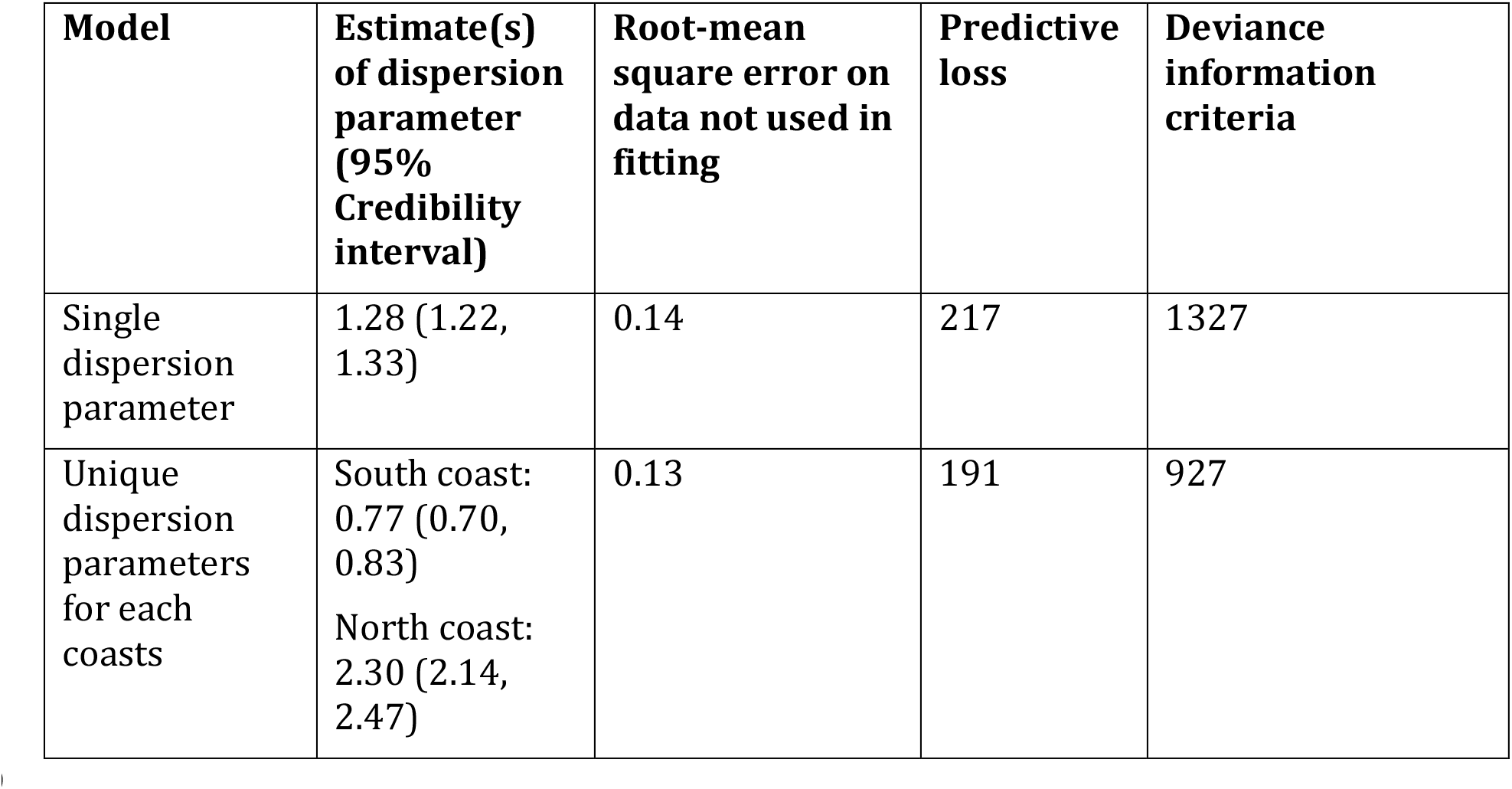
Fit statistics from the two Vanua Levu models. North coast includes Bua Bay

| Model | Estimate(s) of dispersion parameter (95% Credibility interval) | Root-mean square error on data not used in fitting | Predictive loss | Deviance information criteria |
| --- | --- | --- | --- | --- |
| Single dispersion parameter | 1.28 (1.22, 1.33) | 0.14 | 217 | 1327 |
| Unique dispersion parameters for each coasts | South coast: 0.77 (0.70, 0.83)<br>North coast: 2.30 (2.14, 2.47) | 0.13 | 191 | 927 |

The model provided accurate estimates of turbidity. Visualization of modeled turbidity indicated that the large catchments on the north coast contributed to high turbidity in nearby coastal waters (Fig. 2A). On the south-coast the degraded catchments around Savusavu, contributed to moderate levels of turbidity. The moderately sized catchments around western Bua also had a large influence on coastal waters. In comparison the catchments with high forest cover on the south-west coast had little influence on south-west coastal waters. There was some spatial bias in predictions of turbidity when compared to satellite measurements (Fig. 2B). Turbidity was underestimated in inshore areas of the far north coast, particularly near the mouth of the Nasauu River.

**Figure 2.**
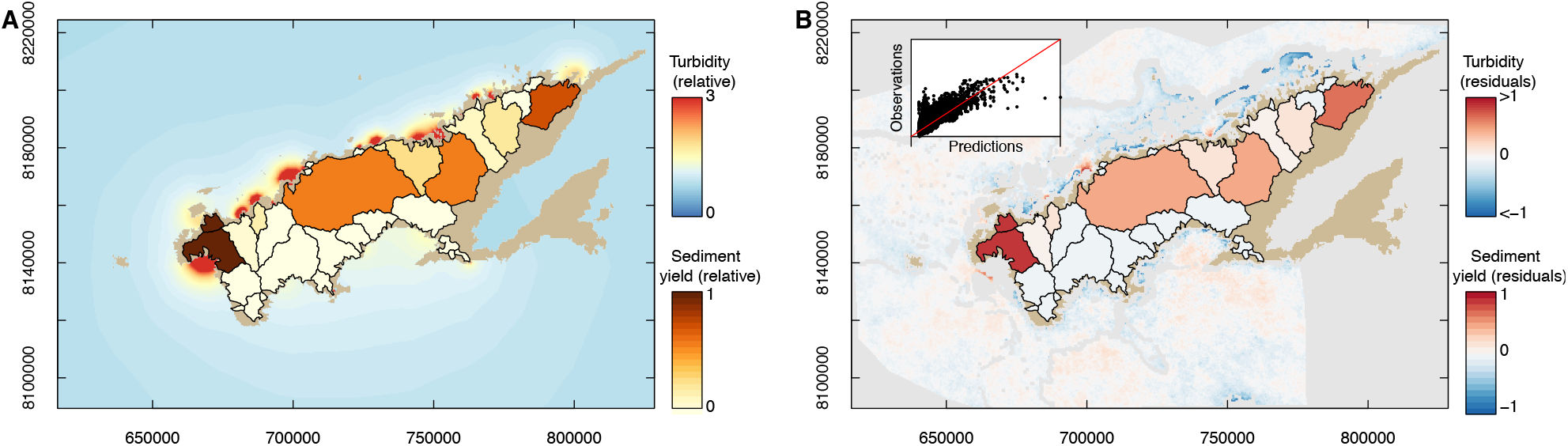
Predicted (A) and residual (B) catchment yields and turbidity values from the model fit for Vanua Levu. The inset on B show a plot of predicted versus residual values. Sediment yields in (A) and residual yields in (B) are scaled relative to the maximum yield value. Residuals for turbidity in (B) are constrained in [−1, 1] to aid visualization, because there were more negative residuals (−4) at the mouth of the Dreketi river.

Estimates of source contributions were generally consistent with the estimates of sediment loadings (Fig. 3), although the estimates for source contributions deviated significantly from the linear relationship with sediment loads for several catchments. In particular, the Bua Bay catchments and Nasauu river and Rukuruku bay catchments were estimated to have a far greater influence on ocean turbidity than the estimates of their sediment loadings suggested.

**Figure 3.**
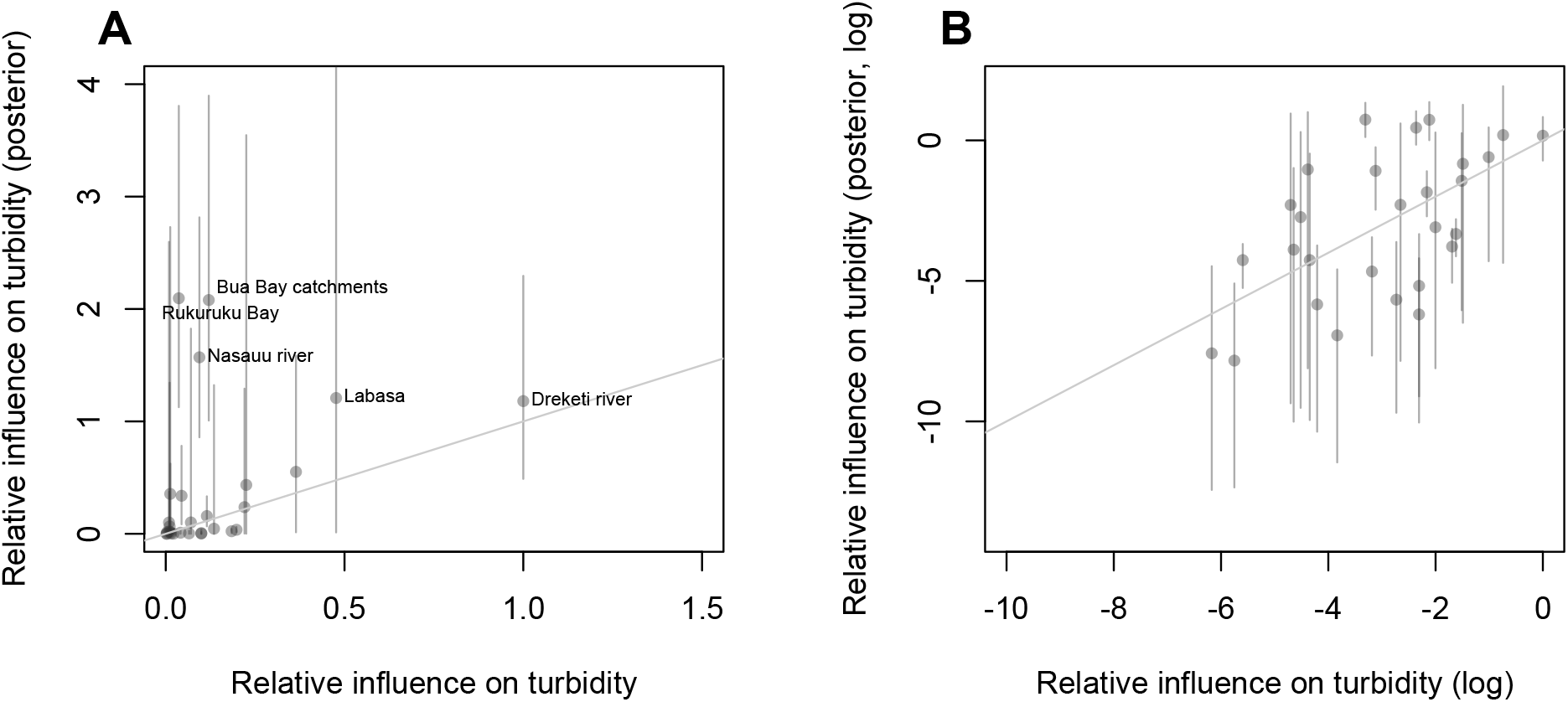
Relative influence of sources on ocean turbidity (A) and logged influence values (B) estimated from the GIS analysis plotted against posterior estimates for relative influence. The x and y axis are scaled to the same units by dividing the x-axis by the slope from the mean estimate of the yield-influence parameter (θ). Error bars show 95% credibility intervals.

The verification analyses relating our predicted turbidity values to observed benthic habitat observations showed some statistically significant relationships, albeit with some uncertainty (Fig. 4). Modeled turbidity was consistent with spatial variation in cover of silt, which was estimated to increase from 2% to 19% from the clearest to most turbid water (Fig 4a, p < 0.05). The cover of sediment sensitive corals declined from an average of 21% to an average of 0.4% with estimated turbidity (Fig 4b, p < 0.05). Algal cover decreased slightly with turbidity, but the relationship was not statistically significant (Fig 4c, p>0.05). Note that change in silt cover and coral cover was greater if sites with extreme turbidity values (>1.3) were removed from analysis.

**Figure 4.**
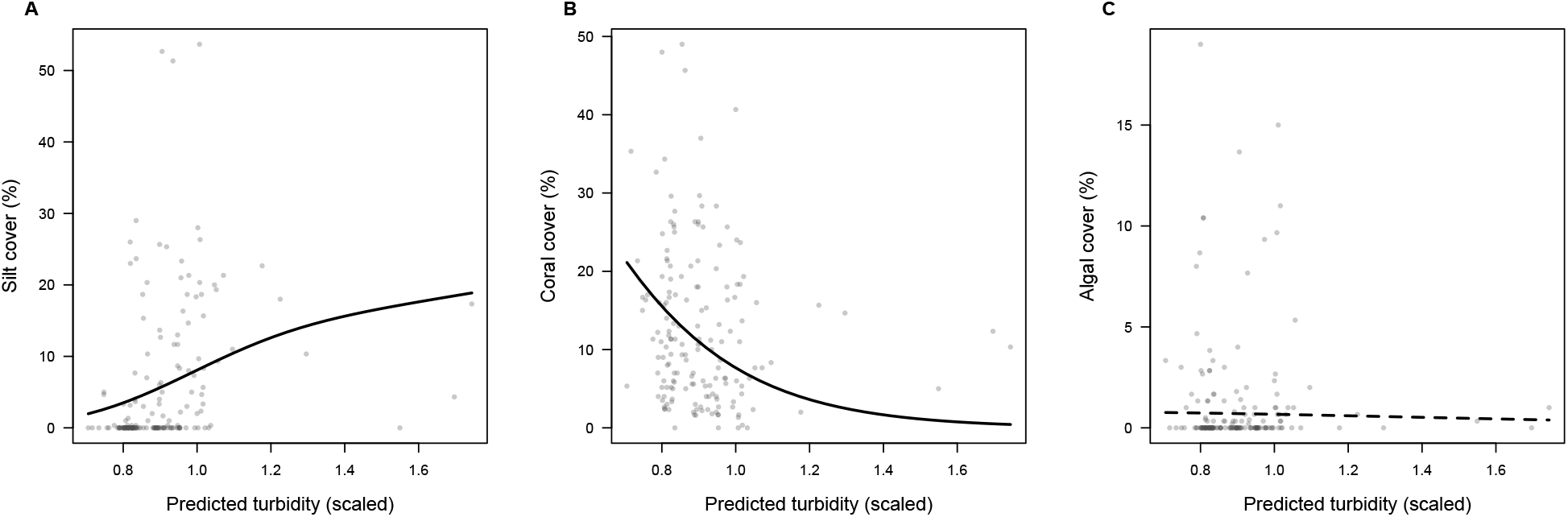
Turbidity estimates from the Bayesian model predict (A) silt cover (zero inflated log-linear model), (B) cover of sediment-sensitive scleractinian coral genera (linear model fit with logit transform) and (C) algal cover (zero inflated logit model). Lines show model fits, where a solid line is statistically significant (p<0.05).

Overall, simulation tests indicated that the model’s estimates of dispersion, source contributions and predictions of water quality had negligible bias across a broad range of parameter and catchment configurations settings (Table 2). The primary cause of bias in estimates was catchment configurations that reduced spatial contrasts in source contributions to water quality. Weak spatial contrasts occurred in two types of data. First, if the observation error on the satellite images was large and the gradient of turbidity from inshore to offshore very weak, the model’s estimates had high bias (>10% difference from the true value) and predictions of turbidity had a large error. Second, if two sources were close together, the contribution of the source with the smaller yield was generally over-estimated, whereas the contribution of the source with the larger yield was underestimated.

**Table 2.** Summary of results from varying parameters for simulating testing of the Bayesian model.

| Test | Bias in estimate of alpha | Variance in estimate of alpha | Bias in source contributions | Variance in source contributions | Root mean square error for water quality predictions |
| --- | --- | --- | --- | --- | --- |
| More negative values of $\alpha$ | <1% for all values | <1% for all values | <1% for all values | <1% for all values | <1% for all values |
| Greater sd obs (sd of $\epsilon_i$) | Negative bias if the sd was large and $\alpha$ low | High variance if the sd was large and $\alpha$ low | Negative bias if the sd was large and $\alpha$ low | High variance if the sd was large and $\alpha$ low | High if the sd was large and $\alpha$ low |
| Sources are closer together | <1% for all values | <1% for all values | Over-predicts smaller source and under-predicts larger source | Higher variance for sources that are closer together |  |
| One source contributes more than the others | <1% for all values | <1% for all values | Bias low, but over-predicts smaller source and under-predicts larger source | Negligible, but increases if sources are close together. | <1% of mean turbidity value for all parameters |

The simulation tests also indicated caution must be taken when interpreting the model’s fit. If rivers differed greatly in their contributions, estimates of water quality were accurate (low root-mean square error), but estimates of source contributions were likely to be biased, particularly if the rivers were close together. Accurate water quality predictions may lead to a false sense of security in extrapolating predictions to other run-off conditions.

## Discussion

Our model provides a rapid and effective tool for estimating the influence of multiple catchments on coastal water quality. The data required to build this model are freely available, making this model useful for regions with limited funding for development of more sophisticated models of pollutant sources that rely on detailed *in situ* data. Further, the model can be rapidly implemented, so can be used to inform development of environmental policy during times of political opportunity ^17^. Model outputs for influence of catchments on water clarity could be used by planners directly, or integrated into simulation models to evaluate different future scenarios of land-use change e.g. ^3,36^. For example, the results from the model for Fiji have provided input into the design of the Bua Province ICM plan for the next 5-10 years. Our modeling was incorporated into the consultation process by informing provincial government and communities which catchments have had the greatest influence on coral reef ecosystems, and therefore need to have sound strategies in place to manage those catchments.

Results indicated that several catchments had a large influence on turbidity. These catchments are some of the most degraded in the region, with native vegetation removed to build towns and grow sugar cane, which has resulted in high erosion rates ^9^. Similar high erosion in other parts of the world has been documented to have substantial impacts on ocean water quality and marine benthic communities ^6,37^. Verification of the model against benthic habitats also demonstrated that turbidity is likely affecting marine benthic habitats, through an increased cover of silt, the stress and eventual loss of sediment-sensitive coral species. Algal cover was not related to turbidity. Multiple processes may drive algal cover, resulting in non-linear response to turbidity, for instance algae may be excluded at low light, and outcompeted by corals in very clear waters. Similarly other studies have also found that algae does not necessarily replace corals on highly turbid reefs ^38,39^. Degradation of fish habitat and turbid waters over coral reefs are of concern to local communities because fisheries and tourism are major sources of livelihoods in the region. Actions reduce deforestation and target catchment restoration in the most degraded catchments may therefore have the greatest benefits for local coastal marine livelihoods.

Our approach offers several advantages over other GIS models of catchment contributions e.g. ^19,20^. Using Bayesian estimation to fit parameter estimates to data enabled us estimate the dispersal of pollutants in the ocean for a given region, rather than using fixed parameters that have been obtained from other regions that may not be locally appropriate. Further, simple GIS models rely on point estimates of catchment contributions to water quality, so do not consider uncertainty in catchment contributions. The Bayesian model estimates uncertainty in catchment contributions. Estimates of uncertainty are useful for decision-makers, because they provide a range over which improvements in catchment land-use are expected to benefit reefs.

There were some discrepancies between the estimates of catchment sediment loadings derived from the GIS analysis and the Bayesian model’s estimates of catchment contributions to turbidity. Further, catchment influence parameters were highly uncertain for several catchments. Such discrepancies may arise due to errors in the satellite measurement of turbidity, processes of erosion that we did not consider in the simple catchment models, or variation in the dispersal of sediment from across different river mouths. Estimates of sediment loadings that were much greater than the GIS estimates may indicate erosion processes we did not consider, such as significant stream bank erosion ^4^ Likewise, where the estimated contributions were much smaller than the estimated sediment loadings, sediment capture and storage processes within hydrological networks may be important ^40^. The discrepancies could also result from oceanographic processes, for instance the effect of some catchments on ocean turbidity may be low if plumes are rapidly dispersed offshore. Where data on catchment processes are available, more detailed process models may provide better estimates of sediment loadings e.g. ^41^. These discrepancies thus indicate key catchments were further empirical work to quantify sediment transport may have the greatest benefit for improving predictions of source contributions to coastal turbidity.

One weakness of our approach is that it does not resolve sub-catchment processes of erosion, so the model can only inform priorities for land-use management at the catchment scale. More detailed catchment models have been used to successfully resolve sub-catchment erosion and thus, can inform on priority areas for restoration within catchments ^42^. However, the most common models for sediment sourcing are based on temperate grasslands, so further work is needed to develop their application to tropical catchments ^8^. In particular, estimates of catchment sediment yield could be improved by accounting for erosion of stream banks, which can be the major source of sediment run-off ^4,43^. The contribution of stream bank erosion to sediment yield could be determined using chemical tracers of sediment sources ^44^ and then catchment scale estimates could then be estimated by mapping streams and remnant riparian vegetation e.g. ^4^. Further development of the model to include stream bank erosion could thus help managers achieve economic development targets for land-use change while avoiding the areas that cause the greatest amount of sediment run-off.

Our model is a simplification of both catchment processes and oceanographic dynamics and several steps could be taken to improve predictions of catchment influences on turbidity at reefs. First, there were no available *in situ* measurements of water quality parameters. Ideally satellite images should be validated against in situ water quality data e.g. ^13,39^ or proxies from coral reef cores e.g. ^45^. To address this, we used satellite products that have previously been validated for other similarly turbid water types ^13^ and verified our estimates against *in situ* data of benthic habitats. Nonetheless, *in situ* water-quality data should be priority for further testing of this model. An advantage of this Bayesian framework is its flexibility to incorporate additional information. For instance, *in situ* measurements of turbidity and sediment concentration could be used as prior information for the estimation of the scaling from sediment loading to turbidity units.

A second caveat is the models of sediment dispersal were simplistic representations of oceanographic processes, including dispersion, tidal transport and wind-driven transport. The power function used to estimate turbidity is a phenomenological representation of sediment dispersion. Thus, bridging the divide between numerical models of sediment dispersion with sophisticated process descriptions e.g. ^46^ and the statistical approach we employed requires further work. The inclusion of additional processes in Bayesian models will require further development to improve the computational efficiency of the estimation algorithm, such as employing customized MCMC algorithms ^23^. One future improvement may be to include the predominant direction of winds in the Bayesian model and allow sediments to disperse further in the direction of winds e.g. ^19^. For instance, we found that model fit was improved considerably with different dispersion parameters for north and south coastlines. The improvement in fit may be due to offshore versus onshore winds on the north versus south coast. More specific parameterizations to account for bathymetric effects on currents may help to resolve spatial auto-correlation in the model’s residuals. Despite the model’s simplistic representation of oceanography, it still explained a large amount of the variance (77%) in the satellite data, suggesting increasing model complexity will provide smaller incremental gains in explanatory power.

We have developed a Bayesian model for estimating the influence of catchments on coastal water quality. The model shows utility for rapidly assessing catchment contributions to water quality in data limited regions and thus may be used to inform planning processes in a timely and cost-effective manner. Building in sediment sourcing models into land-sea planning processes is important to ensure that planners properly account for the downstream effects of actions on land, many of which may impact the livelihoods of coastal people and degrade ecological integrity.

## Acknowledgements

This research was conducted by the Ridges to Reef Fisheries Working Group supported by SNAP: Science for Nature and People Partnership, a collaboration of The Nature Conservancy, the Wildlife Conservation Society and the National Center for Ecological Analysis and Synthesis (NCEAS). CJK was supported by a University of Queensland Research Fellowship. We are grateful to MERIS, the European Space Agency and algorithm developers for making data and algorithms freely available.

